# *C. elegans* harbors pervasive cryptic genetic variation for embryogenesis

**DOI:** 10.1101/008532

**Authors:** Annalise B. Paaby, Amelia G. White, David D. Riccardi, Kristin C. Gunsalus, Fabio Piano, Matthew V. Rockman

**Affiliations:** Department of Biology and Center for Genomics & Systems Biology, New York University, New York, NY 10003, USA; New York University Abu Dhabi, PO Box 129188, Abu Dhabi, United Arab Emirates

## Abstract

Conditionally functional mutations are an important class of natural genetic variation, yet little is known about their prevalence in natural populations or their contribution to disease risk. Here, we describe a vast reserve of cryptic genetic variation, alleles that are normally silent but which affect phenotype when the function of other genes is perturbed, in the gene networks of *C. elegans* embryogenesis. We find evidence that cryptic-effect loci are ubiquitous and segregate at intermediate frequencies in the wild. The cryptic alleles demonstrate low developmental pleiotropy, in that specific, rather than general, perturbations are required to reveal them. Our findings underscore the importance of genetic background in characterizing gene function and provide a model for the expression of conditionally functional effects that may be fundamental in basic mechanisms of trait evolution and the genetic basis of disease susceptibility.

## INTRODUCTION

The nature and extent of functional genetic variation remains a major outstanding question in biology and medicine. For the most part we are in the dark about the degree to which conditions, including genetic and environmental backgrounds, influence a mutation's effect on phenotype. One way to illuminate this problem is to characterize cryptic genetic variation (CGV), the class of mutations that affect phenotype under rare conditions (Gibson & Dworkin 2004; Paaby and Rockman 2014). Unlike mutations that are always silent with respect to phenotype, or mutations that always affect phenotype, CGV is invisible until a perturbation changes the molecular, cellular, or developmental processes that govern its phenotypic expression. For example, CGV may be “released” by environmental exposure, like the modern changes to diet that have been hypothesized to underlie the emergence of complex metabolic diseases in humans (Gibson 2009). Release of CGV may also occur by a genetic change, and inhibition of the heat shock chaperone protein HSP90 has revealed previously-silent mutational effects across many taxa and probably represents a general mechanism that buffers genome-wide functional variation (Queitsh et al. 2002; Yeyati et al. 2007; Jarosz & Lindquist 2010; Chen & Wagner 2012; Rohner et al. 2013).

CGV has long been an important concept in evolutionary biology, principally because it offers a solution to the conundrum of standing genetic variation. If a population is well-adapted to the environment, then it should harbor alleles for the optimal phenotype and not for other, sub-optimal phenotypes. However, well-adapted populations do maintain genetic variation, and sometimes adapt quickly to new environments. One explanation is that some alleles may behave cryptically, silent under established conditions but visible under new conditions (Dobzhansky 1941; Gibson & Dworkin 2004; Paaby & Rockman 2014). Because of its theoretical implications, CGV has been of longstanding interest, and empirical explorations have proved its existence (Waddington 1953; Waddington 1956; Gibson & Hogness 1996; Rutherford & Lindquist 1998; Queitsch et al. 2002; Dworkin et al. 2003; Yeyati et al. 2007; Jarosz & Lindquist 2010; Ledon-Rettig et al. 2010; McGuigan et al. 2011; Duveau & Felix 2012; Cassidy et al. 2013; Rohner et al. 2013). However, as yet some of the most important questions raised by theory remain unanswered (Masel 2006; Masel & Trotter 2010; Gibson & Dworkin 2004; Gibson 2009). How extensive is CGV, under what conditions is it revealed, and how does it influence the genetic architecture of complex traits?

One form of CGV includes modifiers of rare mutations (Paaby & Rockman 2014). The average human genome contains loss-of-function alleles in approximately 300 genes, up to a third of which cause known genetic diseases (Abecasis et al. 2010); disease expression depends on exposure of the disease allele, such as by homozygosity, but also on cryptic variants elsewhere in the genome that act as penetrance modifiers (Hamilton & Yu 2012). In model organisms, this phenomenon is recognized as genetic background effects (Chandler et al. 2013). Our project explores this class of CGV in order to uncover and describe the scope and nature of CGV in a major metazoan process. In *C. elegans*, embryogenesis is robust, and wild-type embryos complete a series of stereotyped cell divisions. Such processes have the potential to accumulate cryptic alleles, as the evolution of redundancy, threshold-mediated traits, and other mechanisms that promote invariant output may shield phenotypes from the effects of some mutations (Gibson & Dworkin 2004). *C. elegans* embryogenesis is also one of the best-described animal processes, following the success of genome-wide screens in identifying embryonic genes, pathways and cellular mechanisms. In this paper, we describe how we leveraged this powerful knowledge base to deliver specific perturbations to the developing embryo, and to uncover a vast reservoir of conditionally functional genetic variation.

## RESULTS

### *C. elegans* harbors substantial CGV for embryogenesis

To test whether wild *C. elegans* populations harbor CGV for embryogenesis, we individually targeted 41 genes in each of 55 wild *C. elegans* strains using RNAi. We targeted maternally-expressed genes with reported embryonic-lethal phenotypes, and excluded from the experiment genes for which we observed developmental or sterility phenotypes in the parental generation. The strains were derived from *C. elegans* populations from around the globe, which on average differ by 8.3 mutations every 10 kb (Andersen et al. 2012). Following gene silencing, the strains exhibited variation in degree of embryonic lethality. The principle of our experimental scheme is that variation in penetrance is due not to allelic differences at the target gene, but to natural genetic variation, otherwise cryptic, elsewhere in the genome.

We used high replication to maximize power for detecting subtle differences in lethality across experimental conditions (Figure 1). Worms were grown in liquid culture in 96-well plates, and RNAi was delivered by feeding the parental generation *E. coli* expressing dsRNA against the target genes (Cipriani & Piano 2011). Each combination of strain and silenced gene was replicated in at least eight wells, and within each well an average of 10 adult worms contributed hundreds of offspring that were screened as dead or alive. Estimates of embryonic lethality were extracted by the image analysis algorithm DevStaR, which was developed to recognize *C. elegans* developmental stages for this specific application (White et al. 2013).

**Figure 1.**
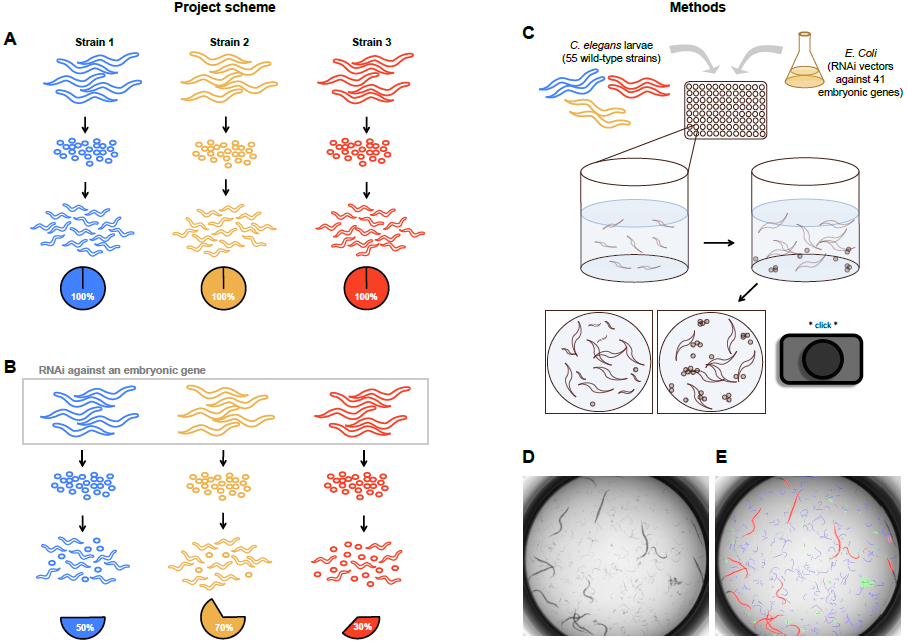
Experimental scheme and methods. (A) Under ordinary conditions, self-fertile hermaphrodites of wild-type *C. elegans* strains lay embryos that all successfully hatch into larvae. (B) In this project, we used RNAi to target maternally-expressed embryonic genes in the germline, inducing embryonic lethality that varied in penetrance across strains, in order to identify cryptic genetic differences in embryonic development. We induced embryonic lethality in 55 wild-type strains by feeding the worms *E. coli* transformed with vectors expressing dsRNA targeting 41 genes. (C) Early-stage (L1) larvae were dispensed into 96-well plates with liquid culture media containing the *E. coli* vectors; worms grew to reproductive maturity and wells were imaged to capture the penetrance of embryonic lethality in the next generation. We replicated each combination of worm strain and RNAi vector in at least eight wells. (D) - (E) Raw images were evaluated using DevStaR (White et al. 2013), a computer algorithm that identifies objects as larvae (blue), dead embryos (green), or adults (red).

From our set of 41 embryonic genes, we further excluded 12 from analysis after examining our results. We excluded two genes that induced growth defects in the parental generation, indicating activity outside of embryogenesis, and another ten genes (Figure 2A) with levels of embryonic lethality indistinguishable from the negative control. We further verified that technical aspects of our experimental design, such as experimental date and the position of wells within a plate, did not confound our ability to detect true differences across the strains. (See the Supplement for details.) To detect CGV that was released after targeting each of the remaining 29 genes, we analyzed variation in embryonic lethality using a generalized linear model with terms for strain, targeted gene, the number of adults per well, experimental date, the strain-by-gene interaction, and pair-wise interactions between the number of adults and both strain and gene (see Materials and Methods). In this model, CGV is represented by the strain-by-gene interaction. This term explains the lethality of a strain under a perturbation after taking into account differences among strains in general sensitivity to perturbations (the strain effect) and differences among genes in the severity of their effects (the targeted gene effect).

**Figure 2.**
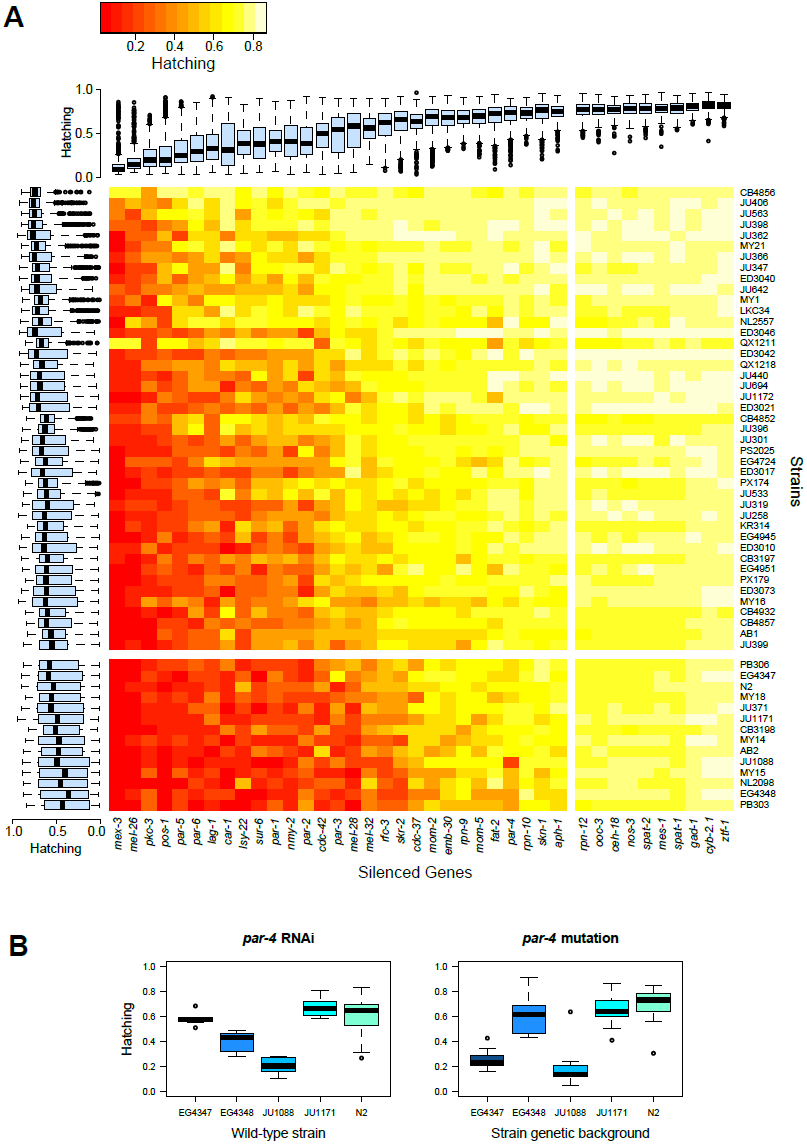
*C. elegans* strains vary in degree of embryonic lethality following gene knock-down. (A) Each cell in the heatmap represents the hatching success for a strain (y-axis) and targeted-gene (x-axis) combination, averaged from at least eight replicate wells. The rows and columns are ordered by average hatching, and boxplots illustrate hatching phenotypes for each strain (across all targeted genes) and for each gene (across all strains). The 14 strains in the lower cluster, which includes 12 wild isolates and the germline RNAi-sensitive controls N2 and NL2098, exhibit lower averages in overall hatching but high variances across genes. Evidence of CGV in these RNAi-sensitive strains can be observed from the raw hatching results plotted here. The right-hand cluster of ten genes with high hatching phenotypes do not show meaningful variation across the strains, and were excluded from downstream analyses. (B) Cryptic genetic variation revealed by targeting *par-4* via RNAi is also revealed by a *par-4* mutation. The temperature-sensitive, EMS-derived mutation *par-4(its57)* was introgressed into four wild strains by backcrossing, and then those strains and the original mutant line, which carries the N2 background, were assayed at 17.5°C. The RNAi results are part of the larger dataset and were collected from liquid culture experiments from a single date batch using RNAi-sensitive strains. The mutation experiments were conducted on solid culture, which may explain the inconsistent hatching phenotype of strain EG4347, as RNAi delivered in liquid versus solid culture can occasionally give divergent outcomes (data not shown).

We observed a highly significant strain-by-gene interaction (F=12.33, DF=1512, p<2x10^−16^), indicating that a substantial fraction of the observed embryonic lethality was associated with the targeted genes in a strain-specific manner. Thus, natural genetic variation between our strains, which is normally cryptic and confers no effect on development at the coarse phenotypic level, affected embryonic survival following knockdown of genes that interact with the cryptic alleles. The strain-by-gene interaction accounted for 15.9% of the total deviance in embryonic lethality (Table 1). The deviance associated with the model term for gene, which represents variation associated with which gene was targeted, accounted for 52.3% of the total deviance. This makes sense, given that our library of RNAi vectors ranged from weakly to strongly lethal. The strain term, which accounts for general strain-wise differences in lethality and likely represents differences in sensitivity to germline RNAi, had a magnitude commensurate with that of CGV, explaining 15.4% of the deviance. Some strains, including the Hawaiian isolate CB4856, are essentially insensitive to germline RNAi delivered by feeding (Tijsterman et al. 2002) and exhibited low lethality across all targeted genes; the majority of the strains showed evidence of intermediate RNAi sensitivity. Although all of the terms in the statistical model were highly significant (Table 1), no additional terms explained more than 2% of the total deviance.

**Table 1.** Factorial analysis of deviance of lethality phenotypes for all 55 wild-type strains, and the subset of 13 RNAi-sensitive strains, in 29 perturbations of germline-expressed genes.

| Analysis on all strains |  |  |  |  |  |  |
| --- | --- | --- | --- | --- | --- | --- |
|  | DF | Deviance | Resid. DF | Resid. Dev | F | p-value |
| NULL |  |  | 17855 | 2201873 |  |  |
| Strain | 54 | 338618 | 17801 | 1863255 | 334.697 | $<10^{-15}$ |
| Silenced gene | 28 | 1152310 | 17773 | 710945 | 2196.584 | $<10^{-15}$ |
| Adults per well | 1 | 35318 | 17772 | 675627 | 1885.113 | $<10^{-15}$ |
| Date | 1 | 2406 | 17771 | 673221 | 128.416 | $<10^{-15}$ |
| Strain x Silenced gene | 1512 | 349415 | 16259 | 323806 | 12.334 | $<10^{-15}$ |
| Strain x Adults per well | 54 | 6715 | 16205 | 317091 | 6.637 | $<10^{-15}$ |
| Silenced gene x Adults per well | 28 | 7358 | 16177 | 309732 | 14.026 | $<10^{-15}$ |
| Analysis on RNAi-sensitive strains |  |  |  |  |  |  |
|  | DF | Deviance | Resid. DF | Resid. Dev | F | p-value |
| NULL |  |  | 5518 | 633652 |  |  |
| Strain | 12 | 14664 | 5506 | 618988 | 73.042 | $<10^{-15}$ |
| Silenced gene | 28 | 463335 | 5478 | 155653 | 989.081 | $<10^{-15}$ |
| Adults per well | 1 | 8332 | 5477 | 147321 | 498.012 | $<10^{-15}$ |
| Date | 1 | 3489 | 5476 | 143832 | 208.532 | $<10^{-15}$ |
| Strain x Silenced gene | 336 | 53123 | 5140 | 90709 | 9.450 | $<10^{-15}$ |
| Strain x Adults per well | 12 | 1947 | 5128 | 88762 | 9.698 | $<10^{-15}$ |
| Silenced gene x Adults per well | 28 | 2736 | 5100 | 86026 | 5.841 | $<10^{-15}$ |

Our ability to estimate the variation in embryonic lethality attributable to CGV, even in the face of incomplete RNAi efficacy, is due to the plurality of embryonic genes we targeted. Specifically, targeting multiple genes increased our power to estimate the strain-by-gene interaction. This approach also permitted us to characterize strain sensitivity to RNAi, and examination of embryonic lethality in an RNAi-sensitive subset of strains further demonstrates extensive CGV. We identified 12 wild isolates with high sensitivity to germline RNAi like the common lab strain N2 (Figure 2A). Together these 13 wild-type strains exhibited similar global levels of embryonic lethality and grouped together in hierarchical clustering according to lethality phenotypes across all targeted genes. Embryonic lethality on individual targeted genes was highly variable within this cluster (Figure 2A). For example, 20% of embryos for strain JU1088 hatched following *par-4* RNAi, compared to 58% of EG4347 embryos. In an analysis of the 13 RNAi-sensitive strains, using the generalized linear model described above, the deviance explained by the strain-by-gene interaction is 3.6 times that of the strain deviance (Table 1). This indicates that differences in general RNAi sensitivity cannot account for the divergent phenotypes we observe, and that *C. elegans* strains harbor substantial functional variation in embryonic gene pathways.

To validate our method of gene silencing by RNAi, we replicated a subset of our results genetically. We introgressed a temperature-sensitive allele of *par-4* into four wild strains and found that the mutation was associated with significant differences in embryonic lethality that were comparable to the RNAi results (Figure 2B). Although one strain showed substantially higher hatching following *par-4* RNAi than with the *par-4* mutation at a semi-restrictive temperature, the near four-fold variation in lethality across the *par-4* introgression strains unequivocally demonstrates the presence of cryptic alleles in the *par-4* network.

We investigated whether the CGV we observed could be due to inter-strain sequence variation at the targeted genes, which might affect the efficacy of RNAi. In the case of *par-4*, our introgression experiment rules out that factor. To examine whether target locus variation might have affected gene silencing in the remainder of our dataset, we looked for polymorphism, relative to the N2 reference sequence from which the RNAi clones were derived, at all targeted-gene loci for 17 strains for which we had whole-genome sequence data. These included 6 RNAi-sensitive strains as well as strain CB4856, which is highly diverged (Andersen et al. 2012) and refractory to germline RNAi delivered by feeding (Tjisterman et al. 2002; Elvin et al. 2011; Pollard & Rockman 2013). Although we observed nucleotide variation in these genes, we observed zero mutations in the exons targeted by the RNAi clones we used. Thus, we exclude RNAi mismatch via target locus sequence variation as a source of the phenotypic variation we observed, based on the complete lack of nucleotide polymorphism at characterized loci and the high efficacy of RNAi even for imperfect sequence matches (Parrish et al. 2000; Rual et al. 2007). We also consider it unlikely that our RNAi vectors silenced unintended targets, as we predicted no off-target sequence matches for the 29 clones used in our final analysis.

### CGV resides in multiple embryonic gene pathways

To identify the sources of variation associated with CGV, we extracted from our statistical analysis the coefficient estimates for the strain-by-gene interaction. The coefficient for each combination of strain and gene provides a quantitative measure of CGV-associated lethality while excluding contributions from the general degree of lethality for a tartgeted gene, the strain effect associated with variation in RNAi sensitivity, the block effect associated with experimental date, and other experimental components of variation.

Each of the 29 genes produced phenotypic variation across the strains that was statistically significant for CGV. That is, an estimated strain-by-gene interaction coefficient was significant (p<0.01) for at least one strain for every gene, which indicates that all silenced-gene perturbations contributed to the highly significant strain-by-gene interaction (Table 1). Most targeted genes were associated with multiple coefficients of high significance (p<0.001). A sufficiently positive (or negative) coefficient indicates that the strain exhibits significantly higher (or lower) hatching on the targted gene than expected, given the strain and gene. We interpret this finding as evidence of pervasive CGV across the multiple pathways represented by the genes we targeted.

However, some genes revealed more CGV than others. Variance in the estimated strain-by-gene interaction coefficients was greatest for genes of intermediate and high lethality (Figure 3). The null expectation is that silenced genes of intermediate lethality will reveal the most variation, since variance in a binomial distribution is highest around a mean of 0.5. This result may support the hypothesis that well-canalized processes, such as those governed by critical genes, can accumulate more CGV under a regime of stabilizing selection.

**Figure 3.**
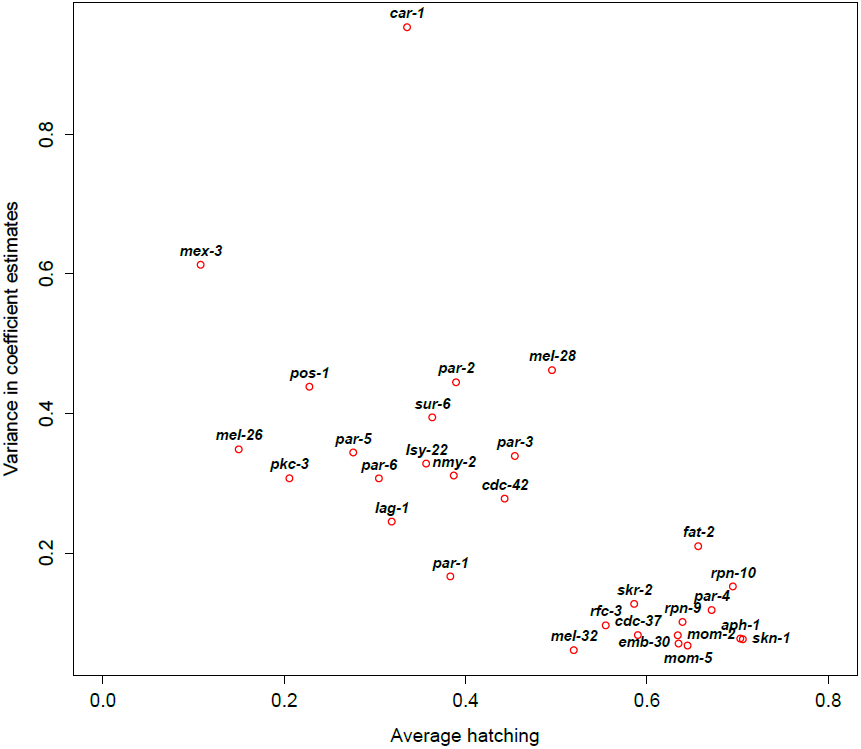
Some targeted genes revealed more CGV than others. The variance in the estimates of the strain-by-gene coefficients, a representation of the degree of CGV revealed by each targeted gene, is plotted here against the average hatching for that gene. Genes with intermediate to high lethality (low hatching) show the greatest variance among strains in coefficient estimates.

### Polarity-associated *par* genes reveal abundant, gene-specific CGV

Targeting the seven genes that *par*tition the cytosol of the early embryo (*par-1* to *-6* and *pkc-3*) and establish anterior-posterior polarity (Kemphues et al. 1988; Hoege & Hyman 2013) revealed abundant CGV. Perhaps surprisingly, the CGV we observed is specific to the silenced gene. For example, strains MY15 and PB306 exhibit low and high hatching, respectively, on *par-3*, but high and low hatching on *pkc-3* (Figures 4, S1). We had hypothesized that if this pathway harbored CGV, we might perturb polarity establishment by silencing multiple pathway genes to uncover the same variation; our hypothesis was motivated by the knowledge that polarity gene products co-localize, interact, and share multiple mutant phenotypes at the cellular level. This is especially true for the three genes whose products comprise the anterior complex (*par-3*, *par-6* and *pkc-3*). Previous work in the wild-type laboratory strain N2 has shown that knock-downs of each of these genes expand PAR-1 and PAR-2 localization in the one-cell embryo, which is normally restricted to the posterior cortex, and mis-orient the AB spindle along the long axis (Etemad-Moghadam et al. 1995; Guo & Kemphues 1995; Tabuse et al. 1998; Hung & Kemphues 1999; Piano et al. 2000; Gonczy & Rose 2005; Cowan & Hyman 2004; Brauchle et al. 2009). In our assay, N2 exhibited consistently low hatching when *par-3*, *par-6* and *pkc-3* were silenced, but other strains showed significant increases in hatching on one or two of the targets (Figure S1). The variability in response indicates that the genetic architecture of polarity-associated CGV includes cryptic variants with low developmental pleiotropy, such that a cryptic allele is conditional on the activity of a specific partner and not the cellular process as a whole. We consider this a phenomenon of low pleiotropy because whatever molecular or cellular effects the cryptic alleles produce, they are not so numerous (or central) that they influence the activity of multiple silenced genes.

**Figure 4.**
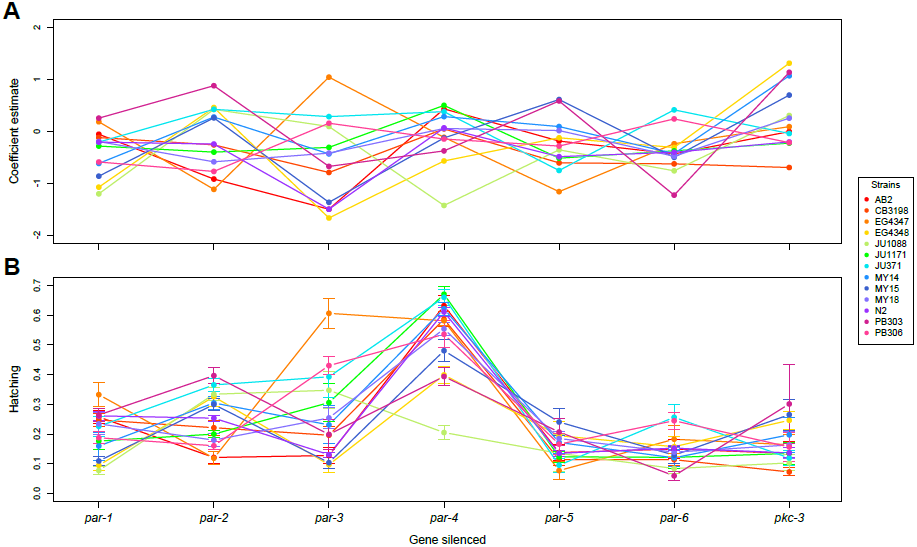
Targeting the 7 genes responsible for partitioning the early embryo reveals substantial CGV. (A) The coefficients for each strain-by-gene combination estimated by the generalized linear model, shown here for the 13 RNAi-sensitive strains, indicate whether a strain demonstrates relatively greater (negative value) or lesser (positive value) lethality for that silenced gene, relative to that expected given the strain and the gene. (B) The degree of embryonic lethality for any given strain is specific to the silenced gene, as shown by the non-parallel lines, and is highly significant across strains. These strains show little variation in the efficacy of RNAi (Table 1), so the coefficients mirror the raw hatching responses. Error bars represent standard error for eight replicates.

### Cryptic phenotypes are likely determined by many loci with common alleles

In order to evaluate the number of loci contributing to the cryptic phenotypes we observed, and to ask whether cryptic alleles are rare or common in populations, we assessed whether genome-wide genetic similarity among strains explained patterns of phenotypic similarity. Using a kinship matrix derived from the SNP genotypes as a measure of strain relatedness, we estimated the genomic heritability of the cryptic phenotypes represented by the strain-by-gene coefficients (Kang et al. 2008). This method estimates heritability associated with alleles of intermediate frequency at many loci, as these are best captured in estimates of strain relatedness.

We found that for most of the lethality phenotypes we measured, many loci contribute to the observed CGV and cryptic alleles are common in natural populations. Of the 29 genes we targeted, 12 induced variation in embryonic lethality for which genomic heritability estimates were greater than 0.80; for 19 genes, estimates were greater than 0.60 (Table 2). However, genomic heritability estimates for *emb-30*, *mel-32*, *mex-3*, *mom-5*, *par-3* and *sur-6* were all 0.00 (Table 2). Because these genes exhibit nonzero variance in their associated strain-by-gene interaction coefficients (Figure 3), including an unusually high variance for *mex-3*, the strains necessarily harbor cryptic genetic differences affecting lethality penetrance. Thus, the genetic architecture of the cryptic variation associated with these genes is likely comprised of few loci, rarer cryptic alleles, or both.

**Table 2.** Genome heritability estimates for CGV phenotypes associated with 29 targeted genes.

| Silenced gene | Heritability estimate | p-value |
| --- | --- | --- |
| *aph-1* | 0.6747 | 0.16 |
| *car-1* | 0.9149 | 0.02 |
| *cdc-37* | 0.7308 | 0.11 |
| *cdc-42* | 0.3639 | 0.29 |
| *emb-30* | 0.0000 | 0.46 |
| *fat-2* | 0.3548 | 0.32 |
| *lag-1* | 0.9075 | 0.01 |
| *lsy-22* | 0.1270 | 0.43 |
| *mel-26* | 0.8245 | 0.05 |
| *mel-28* | 0.8410 | 0.04 |
| *mel-32* | 0.0000 | 0.47 |
| *mex-3* | 0.0000 | 0.76 |
| *mom-2* | 0.7485 | 0.09 |
| *mom-5* | 0.0000 | 0.46 |
| *nmy-2* | 0.4841 | 0.26 |
| *par-1* | 0.7871 | 0.08 |
| *par-2* | 0.9719 | 0.01 |
| *par-3* | 0.0000 | 0.77 |
| *par-4* | 0.9032 | 0.07 |
| *par-5* | 0.6640 | 0.15 |
| *par-6* | 0.9258 | 0.01 |
| *pkc-3* | 0.8136 | 0.06 |
| *pos-1* | 0.7307 | 0.10 |
| *rfc-3* | 0.6958 | 0.13 |
| *rpn-9* | 0.8715 | 0.02 |
| *rpn-10* | 0.8397 | 0.05 |
| *skn-1* | 0.8599 | 0.03 |
| *skr-2* | 0.8961 | 0.02 |
| *sur-6* | 0.0000 | 0.47 |

### Genome-wide associations further support a genetic architecture of low pleiotropy

To explore the genetic architecture of the observed cryptic variation, and to locate genome regions harboring causal variants, we looked for associations between lethality phenotypes and single nucleotide polymorphism (SNP) genotypes across the genome. We performed a separate analysis for each of the 29 targeted genes, and to control for confounding effects we used as phenotypes the strain-by-gene interaction coefficients, which represent strain-wise contributions to phenotypes that are specific to the silenced gene and exclude other components of variation. Of the 29 analyses, nine identified at least one SNP associated with phenotype under a strict Bonferroni-corrected cutoff for significance (Table S1). Across all tests, a total of 19 SNPs or SNP haplotype blocks, defined by SNPs in high linkage disequilibrium (R^2^>0.9), were significant (Table S1).

These associations exceed a strict threshold for significance based on a correction for multiple tests, and likely represent cryptic variants with the largest effect sizes and maximal power for detection. SNPs with p-values exceeding this strict significance threshold might also point to cryptic variants. For example, phenotypes for genes *lsy-22* and *pkc-3* are associated with SNPs on the right arm of chromosome II, with p-values as low as 7.9 × 10^−4^ for *lsy-22* and as low as 1.8x10^−5^ for *pkc-3* (which nevertheless fall short of the cutoff). The SNPs for *lsy-22* are independent of those for *pkc-3* (R^2^=0.03), which reside approximately a megabase away. We introgressed the right arm of chromosome II, containing these sites, from strain N2 into strain EG4348. N2 exhibits low lethality when *lsy-22* is targeted but high lethality on *pkc-3*; EG4348 shows the opposite. In both comparisons, the introgression rescued the original N2 phenotype but had no effect on lethality associated with control gene *mel-32* (Figure 5), for which no SNPs on chromosome II associate with phenotype. These results demonstrate that cryptic variants within the introgression mediate embryonic responses when *lsy-22* and *pkc-3* are targeted, and they suggest that many other SNPs with p-values near the significance threshold may point to true causal loci.

**Figure 5.**
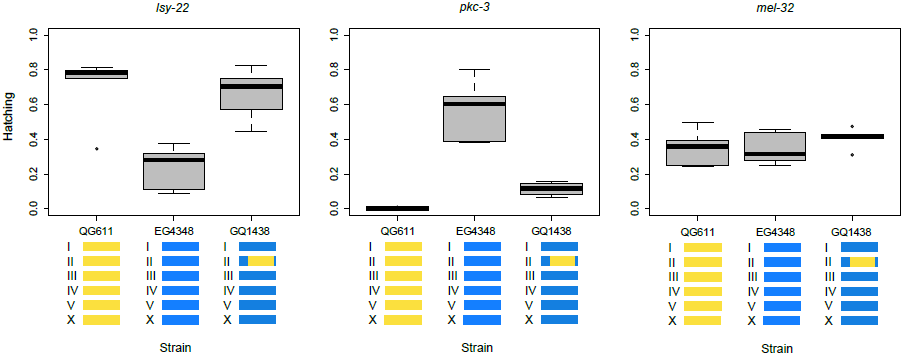
Genetic crosses validate genome-wide association results. Introgression of a section of chromosome II from an N2 strain (strain QG611, in yellow, which carries visible markers in flanking the region of interest) into strain EG4348 (blue) rescues the N2 phenotype on *lsy-22* (F=12.15, DF=2, p=0.001) and *pkc-3* (F=55.87, DF=2, p<0.001) but has no effect on phenotype for control gene *mel-32* (F=0.80, DF=2, p=0.47). Genome-wide analyses found associations between this region and hatching phenotypes for *lsy-22* and *pkc-3*, but not for *mel-32*. The introgression represents approximately 10% of the genome.

We expanded the list of SNPs under consideration to include those with p-values less than 10^−4^, yielding 129 SNPs (in 27 haplotype blocks) associated with CGV (Table S1). These associations were spread across 15 targeted-gene phenotypes. No SNPs lie within or near the locus of the targeted gene, with the exception of one SNP within the *mel-28* locus that associates with the *mel-28* phenotype. This SNP is part of a 6 MB haplotype comprised of 5 significant SNPs, and the *mel-28* phenotype is associated with multiple other SNPs and haplotypes elsewhere in the genome.

Few SNPs associated with more than one phenotype. The lack of shared associations is reinforced by generally low correlations between strain-by-gene coefficients across silenced genes and little to no enrichment across association test p-values. These findings demonstrate that *C. elegans* populations do not appear to harbor cryptic variants that reside in central nodes in gene networks for embryogenesis. If they did, presumably silenced genes that mediate related processes would uncover the same cryptic variants. For example, lethality phenotypes for 4 of the 7 silenced polarity genes (*par-2*, *-4*, *-6* and *pkc-3*) were associated with SNPs, but none were shared. The discrete nature of the genotype-phenotype associations further supports low developmental pleiotropy in the architecture of cryptic genetic variation. In other words, our earlier observation of low pleiotropy, illustrated by perturbation-specific phenotypes for *par* pathway members (Figure 4), is additionally supported by our mapping results, which identify few regions of the genome that appear to affect more than one silenced-gene phenotype.

### Evidence of CGV implicates new network relationships

Although few SNPs were associated with more than one silenced-gene phenotype, the rare instances of multi-gene associations implicate a relationship between those genes (Table S1). In one case, two genes with a known relationship co-associated with SNPs. Both *rpn-9* and *rpn-10* encode non-ATPase regulatory subunits of the proteasome and are predicted to interact with each other (Zhong & Sternberg 2006; Lee et al. 2008); silencing these genes yielded lethality phenotypes associated with two SNPs in a haplotype block on chromosome IV. The haplotype, which spans approximately 10 kb, was also significantly associated with lethality phenotypes for *car-1*, *mom-5*, and *skn-1*, suggesting that a cryptic variant in this region may have pleiotropic effects across multiple embryonic pathways, or that these genes may function together in unknown ways. Phenotypes for *pkc-3*, which encodes an atypical protein kinase and establishes anterior-posterior polarity in the early embryo, and *rfc-3*, a protein-coding gene with homology to DNA replication factors C, are both associated with a SNP on chromosome III. This suggests a potential relationship between *pkc-3* and *rfc-3*, but these genes have no reported interactions or shared functions. Furthermore, their phenotypes are also associated with SNPs on chromosome X. Since the co-association of *pkc-3* and *rfc-3* phenotypes occurs across independent SNPs on different chromosomes (R^2^=0.26), it indicates that the associations are not driven by physical linkage to just one functional variant, but to several, and implicates shared function.

### *C. elegans* strains exhibit differences in the ability to respond to RNAi

Because we measured responses to RNAi against many germline-expressed genes, our results also uncover genetic differences in sensitivity to germline RNAi. Most strains exhibited moderately reduced lethality penetrance relative to the RNAi-sensitive laboratory strain N2, but two strains, the germline RNAi-insensitive strain CB4856 (Tijsterman et al. 2002) and the recently discovered strain QX1211, showed consistently weak lethality penetrance across the targeted genes (Figure 2). In addition to the wild isolates, we also phenotyped the N2 mutant NL2557, which harbors a deletion at *ppw-1* that confers resistance to germline RNAi (Tijsterman et al. 2002). We observed reduced lethality for NL2557 across almost all targeted genes, but unlike CB4856, whose resistance to germline RNAi is largely explained by *ppw-1* function (Tijsterman et al. 2002), NL2557 embryos were dead on genes *mex-3* and *pos-1*. This suggests that CB4856 harbors alleles affecting germline RNAi sensitivity in addition to its nonfunctional copy of *ppw-1*, and is consistent with other work describing germline RNAi resistance in CB4856 as genetically complex (Elvin et al. 2011; Pollard & Rockman 2013).

Our results also uncover genetic variation in sensitivity to somatic RNAi. In addition to the vectors targeting the germline genes, we also included a vector targeting *tubulin* (*tba-2*), which is expressed ubiquitously. All but five strains showed complete sensitivity to somatic RNAi, indicated by developmental arrest on *tba-2*. One of the five exceptions is the N2 mutant NL2098, which is sensitive to RNAi against genes expressed in the germline but carries a deletion at *rrf-1* that confers resistance to RNAi in somatic tissues (Yigit et al. 2006). NL2098 worms reached reproductive maturity but laid dead embryos, indicating that germline RNAi successfully silenced *tba-2* in embryonic development. The other exceptions were wild isolates KR314, JU396, CB4852 and ED3040. These strains also exhibited embryonic lethality on *tba-2* and other germline-expressed genes, but not as completely as NL2098. Thus, these strains must carry variants that affect both germline and somatic RNAi. The strong similarity in phenotypic profiles between N2 and NL2098, which differ only at the *rrf-1* locus, indicates that in general our RNAi vectors target germline-expressed genes and that observed phenotypic differences arise from dynamics in early development. The exceptions to this rule are genes *gpb-1* and *lin-5*; we excluded these from our main analysis of CGV because a few strains failed to develop to reproductive maturity, indicating somatic effects. That these effects were variable across strains indicates either additional variation the machinery for somatic RNAi, or variation in the role of *gpb-1* and *lin-5* in somatic development across strains.

To identify genome regions that mediate germline RNAi, we tested for associations between SNP genotypes and the strain coefficients estimated by the statistical model. No SNP demonstrated significance exceeding a strict Bonferroni correction for multiple tests, but several sets of linked SNPs were suggestive. We examined the polymorphisms identified across 40 wild strains sequenced by the Million Mutation Project (MMP; Thompson et al. 2013) for mutations that are candidates to functionally alter RNAi response and that resided near SNPs with suggestive associations in our genome-wide test. Our top hit was a set of 11 SNPs in perfect linkage disequilibrium with each other, spanning the far left arm of chromosome V (positions 976,982-1,413,337, p=6.09x10^−5^); *ergo-1*, which encodes an Argonaute protein (Yigit et al. 2006), resides within this region and harbors multiple missense mutations and an exonic indel in natural populations (Thompson et al. 2013). Of the 27 strains shared between the MMP and our study, all seven of the strains with the alternate SNP-haplotype at positions V 976,982-1,413,337 also carried at least one protein-coding mutation at *ergo-1*; in contrast, only three of the 20 strains with the reference SNP-haplotype carried a mutation at *ergo-1*. These findings suggest that naturally-occurring mutations at *ergo-1*, if they affect RNAi response, may have driven the association we detected between our measure of RNAi sensitivity and the SNPs on chromosome V. Similarly, two linked SNPs on chromosome III (positions 3,498,885, 3,501,759) were also in association in our analysis (p=6.31x10^−4^); near this region, *nrde-1*, which acts in a nuclear RNAi pathway (Burkhart et al. 2011), contains missense and 3’ UTR mutations. Of the eight strains with alternate SNP-haplotypes at III 3,498,885-3,501,759, seven carry at least one *nrde-1* mutation, compared to five out of the 19 strains with reference SNP-haplotypes. Whether or not these specific mutations prove to be causal variants, our results demonstrate that wild-type *C. elegans* strains harbor extensive genetic variation for germline RNAi, spanning nearly the entire spectrum of sensitivity to resistance.

## DISCUSSION

We have uncovered pervasive cryptic variation in the gene networks for *C. elegans* embryogenesis, indicating that wild populations harbor natural enhancers and suppressors of critical embryonic genes. The extensive knowledge base that enabled this work was necessarily generated in the controlled genetic background of the laboratory strain N2. The limitation of this body of research is that outcomes in a single genetic background represent only a sliver of the high-dimensional reality of gene-gene and gene-environment interactions (Chandler et al. 2013), and our results show that phenotypes of highly conserved genes with critical function can vary widely across genetic backgrounds. This finding contributes to a growing emphasis on the importance of considering genetic background in genetic analysis, and accords with recent work illustrating that even the expression of modifier effects, which represent the relationship between two independent genetic elements, are strongly influenced by variation at other loci (Chari & Dworkin 2013).

To observe CGV we knocked down specific genes, conditions that are likely infrequent in the natural *C. elegans* environment. However, the variation we observed represents heritable differences in developmental mechanism that may affect traits visible to natural selection. *In vitro* experiments have demonstrated that molecules that accumulate cryptic mutations under stabilizing conditions exhibit superior adaptive potential to new molecular substrates (Bloom et al. 2006; Hayden et al. 2011), and cryptic alleles that are partially shielded from selection may be enriched for beneficial mutations that can promote adaptation in future populations (Masel 2006). The degree to which the CGV we observed affects current fitness or the adaptive potential of future populations is as yet unknown, but identification of specific cryptic variants will enable us to test their effects across a diversity of conditions and evaluate their potential role in other aspects of biology and evolution (Duveau & Felix 2012). The CGV may inform change over long time scales; CGV likely precedes the evolution of developmental system drift, a phenomenon where different species show similarity in development but for which the underlying genetic mechanisms are diverged (True & Haag 2001, Felix 2007; Lin et al. 2008; Beadell et al. 2011; Liu et al. 2012; Barriere et al. 2011; Barriere et al. 2012; Duveau & Felix 2012; Verster et al. 2014), as well as the evolution of development itself (Brauchle et al. 2009).

The wild isolates we surveyed varied in sensitivity to germline RNAi, and this variation has the potential to confound our inferences about the nature of the observed CGV. We identified variation in embryonic lethality attributable to genetic differences in germline RNAi by quantifying strain-wise differences in phenotype across all targeted genes. This estimate assumes that strain-wise differences in RNAi efficacy affect germline targets similarly. As yet there is no evidence that germline RNAi, which in our study and others (Tjisterman et al. 2002; Pollard & Rockman 2013) varies dramatically across wild-type genotypes, targets specific genes by distinct mechanisms. Environmentally delivered RNAi is made possible in *C. elegans* by mechanisms that promote RNAi spreading across tissues, leading to non-cell autonomous responses and the experimental expediency of inducing RNAi by feeding. Two classes of RNA spreading-defective mutants have been discovered: those that inhibit RNAi in all tissues, and those that inhibit RNAi in the germline but do not affect somatic RNAi (Whangbo & Hunter 2008). Given that our observations of CGV follow from RNAi perturbations that target a narrow spatio-temporal window, we conclude that genetic differences in mechanisms of RNAi spreading are unlikely to confound our results in a gene-specific way. However, epigenetic processes that interfere with eventual gene activity are perhaps more likely to vary across genotypes. Such processes would include not only RNAi, but any process that depends on small silencing RNAs, including RNA-induced epigenetic silencing (RNAe) or RNA-induced epigenetic gene activation (RNAa) (Ghildiyal & Zamore 2009; Conine et al. 2013; Seth et al. 2013; Wedeles et al. 2013). To date there is no evidence for allele-specific variation in gene silencing or licensing mechanisms, which involve populations of small RNAs derived from the target locus itself, but the question remains entirely unexplored. If the observed strain-by-gene interaction effects we observe are due in part to target-specific RNAi sensitivity mediated by epigenetic variation (a scenario we formally excluded in the case of *par-4*), that epigenetic variation is itself influenced in *trans* by cryptic genetic variation, as shown in our analysis of genomic heritability and GWAS..

Canalized processes have been hypothesized to promote accumulation of CGV, given that mechanisms that stabilize phenotypic output might shield mutational effects (Gibson & Dworkin 2004). Two aspects of our results suggest that the CGV we observe may represent a class of mutations that have escaped elimination (by selection, over evolutionary time) due to their sheltered position with respect to phenotype. First, silencing the most lethal genes revealed the most CGV. If essential genes reside in the most canalized pathways, then canalization may explain the observed CGV. Second, the CGV exhibited low developmental pleiotropy (*sensu* Paaby & Rockman 2013), in that cryptic alleles typically affected lethality phenotypes for only one silenced gene. We hypothesize that this is because alleles that escape purifying selection and persist in a population will be those with neutral or beneficial effects under most conditions; highly pleiotropic alleles that affect multiple aspects of development are more likely to be penetrant, and consequently deleterious. However, we note that our estimates of CGV restricted to the strain-by-gene interaction do not capture any differences in global vulnerability to early embryonic perturbation. Global differences would manifest as strain effects, which we ascribe to variation in sensitivity to germline RNAi. If present, such global vulnerabilities would mean that the strain-by-gene interaction underestimates the scope of CGV and fails to detect instances of widespread pleiotropy. Low pleiotropy nevertheless remains a central observation in our study, as lack of pleiotropy can be observed directly from the raw phenotype estimates in RNAi-sensitive strains, including those following perturbation of *par* pathway members.

Some of our findings may be particular to our system. *C. elegans* persist primarily as selfing hermaphrodites in the wild and lineages exhibit shared haplotypes defined by extensive linkage disequilibrium across the genome (Andersen et al. 2012). Such conditions should promote the evolution of background-specific fitness effects given stabilizing selection on phenotype, which is exactly what we see in our observations of pervasive CGV. *C. elegans* populations are also highly inbred, which means that cryptic alleles rarely segregate as hidden recessives as they might in outbred populations with greater heterozygosity. The CGV we characterize here represents mutations that have accumulated, following exposure in homozygous genotypes, in lineages with occasional recombination. In this system, recombination and the subsequent creation of new haplotype combinations likely represents an important source of perturbation for revealing CGV (Masel & Trotter 2010).

The CGV we discovered opens the door to fresh investigation into the developmental genetics of embryogenesis. The observations of CGV associated with the seven major polarity genes, *par-1* to *par-6* and *pkc-3*, provide a particularly powerful opportunity for improving our resolution of developmental processes. In particular, the shared function of *par-3*, *par-6* and *pkc-3*, which comprise the anterior complex (Etemad-Moghadam et al. 1995; Guo & Kemphues 1995; Tabuse et al. 1998; Hung & Kemphues 1999; Piano et al. 2000; Gonczy & Rose 2005; Cowan & Hyman 2004; Brauchle et al. 2009) contrasts with our observation that silencing these genes induces high variability in embryonic lethality across genetically diverse strains. Examination of spindle orientation and PAR-1 and PAR-2 localization in different strains will identify which of these aspects of early cell division are associated with CGV, which are more and less tolerant of phenotypic variation, and how that relates to the known activity of polarity genes. In its entirety, our embryonic lethality dataset identifies many more strain and gene combinations for which to examine cellular phenotypes. By leveraging existing knowledge of embryonic gene mechanisms and phenotypes, such evaluations of natural genetic variation will provide valuable and complementary insight into the shape and flexibility of embryonic gene networks.

## MATERIALS AND METHODS

### C. elegans *strains*

We evaluated laboratory strain N2 and 54 wild-type strains derived from populations around the world. Strains and their provenances are detailed in the Supplement. We also assayed N2 mutants NL2557, which carries a deletion at *ppw-1* (*pk1425*) that confers resistance to RNAi in the germline (Tijsterman et al. 2002), and NL2098, which carries a deletion at *rrf-1* (*pk1417*) that confers resistance to RNAi in somatic tissues (Yigit et al. 2006). These were provided by the Caenorhabditis Genetics Center, which is funded by NIH Office of Research Infrastructure Programs (P40 OD010440).

#### Phenotyping embryonic lethality in liquid culture

Worms were grown to large numbers on agarose-media plates, and healthy embryos at least two generations past starvation or thawing were collected using standard bleaching techniques. For each strain, ~10,000 embryos were plated onto a 15 cm agarose-media plate densely seeded with *E. coli* OP50. Worms were reared at 20°C with food until gravid, then bleached and the embryos synchronized to the arrested L1 larval stage by rocking in M9 buffer overnight at 20°C. Following the high-throughput phenotyping methodology described in Cipriani & Piano (2011), larvae were washed and diluted to ten worms per 20 uL of S buffer with additives. Worms were dispensed with a peristaltic pump (Matrix Wellmate) in 20 uL volumes into wells of flat-bottomed 96-well plates (in rows, 8 strains per plate) already containing 30 uL of the appropriate RNAi feeding bacteria. Each plate was replicated eight times, and N2 was dispensed on every plate. After dispensing, plates were stored at 20°C in sealed humid chambers for five days. Three sets of eight worm strains were dispensed per experimental cycle; we performed a total of three cycles over three months.

#### RNAi vectors

We silenced 41 germline-expressed genes and one somatic gene (*tba-2*) by feeding the worms HT115 *E. coli* bacteria expressing dsRNA for their targets. Bacteria had been transformed with L4440 RNAi feeding vectors into which target DNA had been cloned (Timmons et al. 2001) and carried genes for ampicillin and tetracycline resistance. We also included *E. coli* carrying the empty L4440 vector, for a total of 43 RNAi vectors in the survey. The list of RNAi vectors and their sources are detailed in the Supplement; the genes we targeted (excluding *gpb-1*, *lin-5*, and *tba-2*, which exhibited somatic effects) are shown in Figure 2.

We constructed a frozen RNAi bacterial feeding library in 96-well plates with 20% glycerol. The bacteria were distributed across the plates in columns (12 vectors per plate); the *mom-2* vector was included on every plate. Using a 96-pin replicator, bacterial colonies were transferred from the frozen libraries and grown on LB agar plates (100 ug/mL ampicillin, 12 ug/mL tetracyclin). LB broth (50 ug/mL ampicillin) in 96-deep-well plates was inoculated from the solid cultures using the pin replicator and grown overnight in a 37°C shaker. Cultures were induced with 1 mM IPTG for two hours and dispensed into 96-well flat-bottom plates using a Tecan Aquarius robot.

Off-target predictions for our RNAi clones were generated from a sliding window analysis of matching 21-mers between the RNAi reagent and the *C. elegans* reference genome (ce10).

#### Image acquisition and data extraction

Five days after the experimental cycle was initiated, the L1 larvae had developed into egg-laying adults and consumed the RNAi bacteria so that the wells were optically clear. Wells were photographed at the point at which viable embryos had hatched but not developed past early larval stages. We captured single images of each well using a DFC340 FX camera and a Z16 dissecting microscope (Leica Microsystems, Inc.), a Bio-precision motorized stage with adaptors for the 96-well plates and stage fittings (Ludl, Inc.), and Surveyor software from Media Cybernetics, Inc. We used a 1.2 ms exposure at 17.3X magnification.

Data were extracted from the images using the automated image analysis system DevStaR (White et al. 2013). DevStaR is an object recognition machine that classifies each object in the image as an adult, larva or embryo using a support vector machine and global shape recognition. Embryonic lethality estimates were derived from the proportion of embryos in each well relative to all progeny (embryos plus larvae). During the development of DevStaR, each of the approximately 30,000 images in this experiment were manually evaluated and assigned qualitative scores for the number of embryos and the number of larvae, and exact counts were determined for the adults in each well. These data provided independent phenotype estimates and demonstrate that DevStaR is accurate and reliable (White et al. 2013), and we used the manually-collected adult count data in our analyses evaluating the number of adults in each well.

#### Statistical analyses

The counts of dead embryos and living larvae from each experimental well were bound together as a single response variable and modeled using a generalized linear model with a quasi-binomial error structure. In the central analysis, in which we evaluated 55 strains and 29 genes, the model included main effects of strain, silenced gene, number of adult worms per well, and experimental date; and interaction terms for strain-by-gene, strain-by-adults and gene-by-adults, in the form:

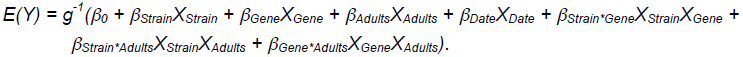

where *g^−1^* represents a logit link function. The analysis was conducted using the *glm* function in R (R Development Core Team 2010) and model fit was examined with the deviance statistic. Coefficients from the strain-by-gene interaction term in this model were used as estimates of CGV. The significance of each coefficient was computed by assessing the coefficient ratio against the *t*-distribution using the *summary.glm* function. We also performed a mixed-model analysis using the *glmer* function in the R package *lme4* (Bates et al. 2011), with a logit link function and a binomial error structure, in which all effects except the number of adults were specified as random. Results from this analysis were consistent with the fixed-effects analysis, including tight correlation between the fixed-effect coefficients and the mixed-effect estimates and between the downstream GWAS results; we only report results from the fixed-effects analysis. Other analyses, including those exploring confounding effects of experimental design, fitted models with additional terms for well position and bacterial source to subsets of the data. To identify best-fitting models, terms were sequentially reduced from the full model and model comparison was achieved with the F statistic.

#### Genome-wide association tests

Association analyses of the gene-specific embryonic lethality phenotypes were implemented with the *emma.ML.LRT* function in the R package *emma*, which controls for population structure using a kinship matrix and performs efficient mixed-model association using maximum likelihood (Kang et al. 2008). The kinship matrix was determined from a total of 41,188 SNPs across 53 strains; we excluded strains CB4856 and QX1211, as they are essentially insensitive to RNAi in the germline. The SNP genotypes are as described in Andersen et al. (2012) and were downloaded from the website of E. Andersen (http://groups.molbiosci.northwestern.edu/andersen/Data.html). We assayed six wild isolates not fully genotyped by that study; in the Supplement we describe our imputation method for assuming their genotypes. The phenotype values were the coefficients estimated from the strain-by-gene interaction by the generalized linear model, as they include strain contributions to lethality minus the strain effect of RNAi sensitivity, the date effect, and other effects of experimental design. We evaluated SNPs with minor allele counts of 6 or more, which allowed us to interrogate 9,362 SNPs. Of these, 3,057 exhibit unique genotype identities across the 53 strains, and the strict threshold for significance, following Bonferroni correction for multiple tests, was determined at 0.05/3,057, or 1.6x10^−5^. Genomic heritability estimates for each of the cryptic phenotypes represented by the strain-by-gene coefficients was determined from the genetic and residual error variance components estimated by restricted maximum likelihood, using the function *emma.REMLE*. Significance was tested by 1,000 permutations of strain phenotypes.

The association analysis for RNAi sensitivity was performed as above, using as phenotypes the strain coefficients estimated by the generalized linear model. We included 54 strains, excluding only CB4856 as its insensitivity to germline RNAi is largely explained by *ppw-1* (Tjisterman et al. 2002; Pollard & Rockman 2013), for which we interrogated 9,690 SNPs.

#### Validation of CGV by mutation and introgression

We placed the temperature-sensitive, embryonic-lethal *par-4* mutation *its57* into different wild-type genetic backgrounds by backcrossing the mutant strain to four wild isolates for six generations. Embryonic lethality was measured at a permissive 17.5°C on solid culture plates and compared to strains that had been treated with RNAi against *par-4* in a single date batch. Similarly, we introgressed a region on chromosome II from the strain N2 into the background of strain EG4348 by backcrossing for 20 generations. The effects of the introgression on embryonic lethality were tested by feeding the strains RNAi against *lsy-22*, *pkc-3* and *mel-32* on solid culture using standard techniques. Details can be found in the Supplement.

#### Genome sequencing

Seventeen strains (AB1, AB2, CB3198, CB4852, CB4856, EG4347, EG4348, JU319, JU371, JU1088, JU1171, MY1, MY16, MY18, PB306, PX174, PX179) were examined for sequence variation at the RNAi target sites. Sequences were derived from 100-bp paired-end reads run on an Illumina HiSeq 2500 that were mapped to the N2 reference (ce10) using *stampy* (Lunter & Goodson 2011) and variant-called with *samtools* (Li et al. 2009).

## AUTHOR CONTRIBUTIONS

The experiments in this project were designed by ABP and MVR. FP and KCG developed the phenotyping pipeline, and AGW, FP and KCG developed the image analysis algorithm DevStaR. DDR performed whole-genome sequencing on the worm strains. ABP conducted the experiments, ABP and MVR analyzed the data, and ABP wrote the paper.

## ACKNOWLEDGMENTS

We thank Miyeko Mana for RNAi clones, Huey-Ling Kao for help in computational data management, Firoz Ahmed for RNAi off-target predictions, and Patricia Cipriani, Eliana Munarriz and Katherine Erickson assistance with the high-throughput phenotyping platform. Kirk Rattanakorn, Caleb Karmel, and Amalia Stavropoulos scored data images, which provided validation for DevStaR. We also thank Luke Noble for assisting in the analysis of whole-genome sequence data and Dan Pollard for useful discussions and advice.

## SUPPLEMENTAL MATERIALS AND METHODS

### C. elegans *strains*

The 55 wild-type strains we evaluated included the common wild-type laboratory strain N2, originally derived from Bristol, England, and 54 strains from other natural populations: AB1, AB2 (Adelaide, Australia), CB3197, PS2025 (Altadena, CA, USA), CB3198 (Pasadena, CA, USA), CB4852 (Rothamsted, England), CB4856 (Hawaii, USA), CB4857 (Claremont, CA, USA), CB4932 (Taunton, England), ED3010, ED3017, ED3021 (Edinburgh, Scotland), ED3040 (Johannesburg, South Africa), ED3042, ED3046 (Western Cape, South Africa), ED3073 (Limuru, Kenya), EG4347, EG4348, EG4945, EG4951 (Salt Lake City, UT, USA), EG4724 (Amares, Portugal), JU1088 (Japan), JU1171, JU1172 (Chile), JU258 (Madeira, Portugal), JU301 (LeBlanc, France), JU319, JU347 (Merlet, France), JU362, JU366, JU371, JU694 (Franconville, France), JU396, JU398, JU399, JU406 (Hermanville, France), JU440 (Beauchene, France), JU533 (Primel, France), JU563 (Sainte Barbe, France), JU642 (Le Perreux, France), KR314 (Vancouver, Canada), LKC34 (Madagascar), MY1 (Lingen, Germany), MY14, MY15, MY16 (Mecklenbeck, Germany), MY18, MY21 (Roxel, Germany), PB303 (isolated from an isopod from Ward's Biological Supply), PB306 (isolated from an isopod from Connecticut Valley Biological Supply), PX174 (Lincoln City, OR, USA), PX179 (Eugene, OR, USA), QX1211 (San Francisco, CA, USA), and QX1218 (Berkeley, CA, USA). Isolates were acquired from the Caenorhabditis Genetics Center or kindly shared by members in the worm community.

#### RNAi vectors

The majority of the RNAi vectors we used were obtained from the Ahringer feeding library (Kamath & Ahringer 2003). These included: *aph-1* (VF36H2L.1), *car-1* (Y18D10A.17), *cdc-37* (W08F4.8), *cdc-42* (R07G3.1), *ceh-18* (ZC64.3), *cyb-2.1* (Y43E12A.1), *emb-30* (F54C8.3), *fat-2* (W02A2.1), *gad-1* (T05H4.14), *lag-1* (K08B4.1), *lin-5* (T09A5.10), *lsy-22* (F27D4.2), *mel-26* (ZK858.4), *mel-28* (C38D4.3), *mel-32* (C05D11.11), *mes-1* (F54F7.5), *mex-3* (F53G12.5), *mom-2* (F38E1.7), *mom-5* (T23D8.1), *nmy-2* (F20G4.3), *nos-3* (Y53C12B.3), *ooc-3* (B0334.11), *par-1* (H39E23.1), *par-2* (F58B6.3), *par-3* (F54E7.3), *par-5* (M117.2), *par-6* (T26E3.3), *pkc-3* (F09E5.1), *pos-1* (F52E1.1), *rfc-3* (C39E9.13), *rpn-10* (B0205.3), *rpn-12* (ZK20.5), *rpn-9* (T06D8.8), *skn-1* (T19E7.2), *skr-2* (F46A9.4), *spat-1* (F57C2.6), *spat-2* (Y48A6B.13), *sur-6* (F26E4.1), *tba-2* (C47B2.3), and *ztf-1* (F54F2.5). We also used two feeding vectors created and kindly shared by M. Mana, for genes *gpb-1* (F13D12.7) and *par-4* (Y59A8B.14).

#### Experimental replication and controls

Because we arranged worm strains in fixed rows and RNAi vectors in fixed columns across the 96-well experimental plates, well position was a potentially confounding source of variation in the data. The source of each bacterial culture was also potentially confounding, as each culture was grown independently for each strain on a plate. To estimate the contribution of these variables to the lethality phenotypes, we examined hatching variation for strain N2 on silenced gene *mom-2*, which we included in every plate. The dataset includes counts of dead and alive offspring from 285 experimental wells. Independent cultures of *E. coli* bacteria expressing dsRNA against *mom-2* only weakly affected hatching (F=3.12, DF=2, p=0.046) (Table S2), and whether a well was on the edge, near the edge, or in the center of the plate had no effect on phenotype (F=1.39, DF=2, p=0.251).

With the exception of N2, strains were assayed in one of three date batches. To evaluate the relative importance of date, we examined the N2 lethality phenotypes for all 29 lethality-inducing genes across the three dates. While the date effect was statistically significant, it explained only 1.9% of the deviance; the gene effect explained 86.6% of the deviance (Table S3). The model that best fits the data also includes main and interaction terms for the number of adults per well, but their effects are similarly negligible.

#### Excluding genes from the analysis

Although we evaluated 41 genes in our experiment, in our final analysis we included results only for 29. Silencing *gpb-1* and *lin-5* induced growth defects in multiple strains such that the parental generation of worms failed to develop to reproductive maturity, indicating that these genes have effects outside of embryogenesis. We also identified ten genes (Figure 2) that induced no or extremely low embryonic lethality. Non-zero embryonic lethality estimates were extracted from the experimental wells in which these genes were silenced (Figure 2), but the estimates are slightly inflated. In wells in which all or nearly all embryos hatch into larvae, discarded eggshells coalesce in the crowded well and can be erroneously identified as a clump of dead embryos by the image analysis algorithm. To test whether the slight differences in lethality estimates across strains were nevertheless meaningful for any of the low-lethality genes, we analyzed the data from all 41 embryonic genes as well as the RNAi empty vector (negative control) using the full statistical model. We then computed the variance in the estimated strain-by-gene coefficients across all strains for each silenced gene. As the coefficients include the contribution to embryonic lethality that is strain-specific for a silenced gene, a high coefficient variance indicates that strains differ meaningfully in response to the silenced gene treatment. However, the variances in strain coefficients were universally low for all of the low lethality genes, equal to or less than the variance in strain coefficients for the empty vector (on which all strains hatch completely). Consequently, we concluded that results from the ten low lethality genes were not meaningful and we excluded them from analysis.

#### Genotype imputation

Six wild isolates in our study were not fully genotyped using the RAD-seq method by Andersen et al. (2012), and we used the following imputation procedure to assume genotypes at the full set of SNPs. If the strain was identical at the 1454 SNPs assayed by Rockman & Kruglyak (2009) to a strain genotyped by RAD-seq, we used the RAD-seq data of the matching strain. This allowed us to use genotype data of CB4854 for CB3197; JU310 for JU301; JU311 for JU319; JU367 for JU371, and MY10 for MY21. In each of these cases, multiple RAD-sequenced strains collapse into groups of strains that are also identical at the 1454 SNPs, suggesting that this procedure is reliable. Only in the case of JU366 do we encounter uncertainty. At the 1454 SNP markers, this strain is identical to JU360, JU363, and JU368 (and three others not RAD-sequenced). JU360 and JU368 have identical RAD-seq haplotypes, but JU363 is different at 224 sites (of which 164 were tested for association with phenotype). We substituted both JU360 and JU363 as proxies for JU366 and ran the full GWAS pipeline twice; the differences in outcome were negligible, with extremely tight correlation among SNP p-values across all tests and no differences in the set of statistically significant SNPs.

#### Validation of CGV by mutation

We introgressed the EMS-derived mutation *par-4(its57)* into five wild-type strains by mating hermaphrodites of the mutant KK300 to wild-type males followed by backcrossing for six generations. The mutation is a single nucleotide change causing a proline to serine substitution, which induces embryonic lethality at higher temperatures (Watts et al. 2000). We conducted the crosses at 14°C and tracked the mutation by PCR and digestion with restriction endonuclease XmnI, which cuts the mutant but not the wild-type. Embryonic lethality was measured by manually counting the number of hatched larvae and dead embryos, laid by 10 hermaphrodites on individual agarose-media plates seeded with OP50 *E. coli*, over three days of sequential transfers, for each introgressed strain. This experiment was performed at 17.5°C. The results were compared to strains that had been treated with RNAi against *par-4* in a single date batch.

#### Validation of CGV by introgression

We created the strain QG611, which carries two markers (*mIs12*, expressing GFP in the pharynx, and *juIs76*, expressing GFP in the motor neurons) in the N2 wild-type background. The markers are positioned at the approximate middle and right end of chromosome II, respectively (precise locations are unknown), which flank the region for which *lsy-22* and *pkc-3* phenotypes were associated. We crossed QG611 to wild-type strain EG4348 and then backcrossed to EG4348 for 20 generations, retaining the N2 introgression by selecting for the double markers. The introgression strain, QG1438, carries the N2 haplotype from approximately II 3,174,000 to the right of II 14,430,751. To test the effect of the introgression on *lsy-22*, *pkc-3* and *mel-32* gene silencing, RNAi was induced by feeding on agarose plates following standard protocols (wormbook.org). Test strains included QG611 (the GFP constructs in QG611 have no effect on phenotype relative to N2, data not shown), EG4348, and QG1438. Test worms were singled onto plates, 6 replicates each, at the L4 (*lsy-22* and *pkc-3*) or L1 (*mel-32*) stage following bleaching and developmental synchronization; worms were transferred daily for three days and the number of dead embryos and hatched larvae were counted 24 hours after transfer. The data were analyzed using the statistical approach described elsewhere; here, the generalized linear model tested the effect of strain.

**Figure S1.**
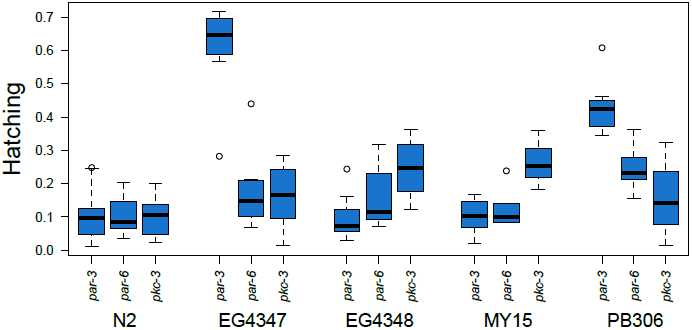
Silencing members of the anterior polarity complex (*par-3*, *par-6* and *pkc-3*) induces severe and consistent embryonic lethality in strain N2, but not all wild-type strains. The raw data plotted here show hatching phenotypes for four wild isolates in addition to N2, and represent results that are typical for all 13 RNAi-sensitive strains. For consistency, only data collected from a single date batch is shown for each strain.

**Table S1.**
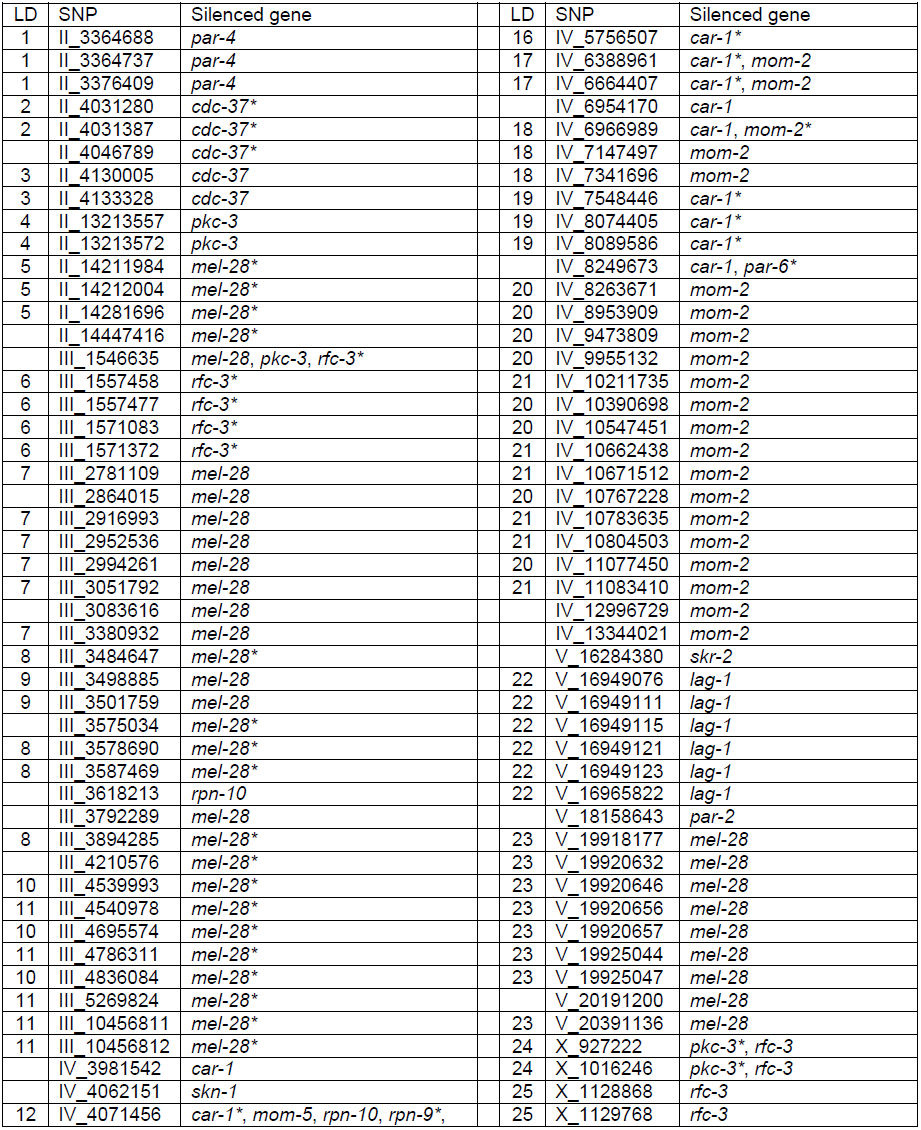
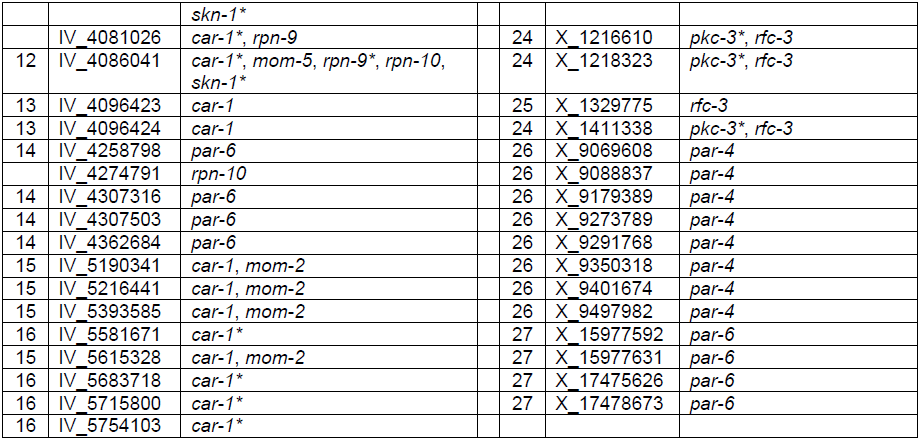
Genome-wide SNPs associated with hatching phenotypes with p-values < 0.0001 and < 1.64x10^−5^ (*). The LD column indicates clusters of SNPs in strong disequilibrium with each other (R^2^ > 0.90) across our test strains.

**Table S2.** Factorial analysis of deviance of strain N2 lethality on silenced gene *mom-2*.

|  | Df | Deviance | Resid. Df | Resid. Dev | F | Pr(>F) |
| --- | --- | --- | --- | --- | --- | --- |
| NULL |  |  | 284 | 9191.7 |  |  |
| Date | 1 | 1060.26 | 283 | 8131.4 | 38.397 | 2.0 x10<sup>-09</sup> |
| Bacterial source | 2 | 172.47 | 281 | 7958.9 | 3.123 | 0.04556 |

**Table S3.** Factorial analysis of deviance of strain N2 lethality phenotypes across 29 silenced genes.

|  | Df | Deviance | Resid. Df | Resid. Dev | F | Pr(>F) |
| --- | --- | --- | --- | --- | --- | --- |
| NULL |  |  | 2254 | 280706 |  |  |
| Silenced gene | 28 | 221081 | 2226 | 59624 | 378.3275 | <2 x10<sup>-16</sup> |
| Date | 1 | 6090 | 2225 | 53534 | 291.8186 | <2 x10<sup>-16</sup> |
| Adults per well | 1 | 249 | 2224 | 53285 | 11.9248 | 0.00056 |
| Silenced gene x Date | 28 | 5265 | 2196 | 48020 | 9.0099 | <2 x10<sup>-16</sup> |
| Silenced gene x Adults per well | 28 | 2423 | 2168 | 45597 | 4.1467 | 2.7x10<sup>-12</sup> |

